# In-depth Bioinformatic Analyses of Human SARS-CoV-2, SARS-CoV, MERS-CoV, and Other *Nidovirales* Suggest Important Roles of Noncanonical Nucleic Acid Structures in Their Lifecycles

**DOI:** 10.1101/2020.04.09.031252

**Authors:** Martin Bartas, Václav Brázda, Natália Bohálová, Alessio Cantara, Adriana Volná, Tereza Stachurová, Kateřina Malachová, Eva B. Jagelská, Otília Porubiaková, Jiří Červeň, Petr Pečinka

## Abstract

Noncanonical nucleic acid structures play important roles in the regulation of molecular processes. Considering the importance of the ongoing coronavirus crisis, we decided to evaluate genomes of all coronaviruses sequenced to date (stated more broadly, the order *Nidovirales*) to determine if they contain noncanonical nucleic acid structures. We discovered much evidence of putative G-quadruplex sites and even much more of inverted repeats (IRs) loci, which in fact are ubiquitous along the whole genomic sequence and indicate a possible mechanism for genomic RNA packaging. The most notable enrichment of IRs was found inside 5′UTR for IRs of size 12+ nucleotides, and the most notable enrichment of putative quadruplex sites (PQSs) was located before 3′UTR, inside 5′UTR, and before mRNA. This indicates crucial regulatory roles for both IRs and PQSs. Moreover, we found multiple G-quadruplex binding motifs in human proteins having potential for binding of SARS-CoV-2 RNA. Noncanonical nucleic acids structures in *Nidovirales* and in novel SARS-CoV-2 are therefore promising druggable structures that can be targeted and utilized in the future.

## 1 Introduction

The order *Nidovirales* is a monophyletic group of animal RNA viruses. This group can be divided into the six distinct families of *Arteriviridae, Coronaviridae, Mesnidovirineae, Mononiviridae, Ronidovirineae* and *Tobaniviridae*. All known *Nidovirales* have single-stranded, polycistronic RNA genomes of positive polarity (Modrow et al., 2013). Due to the Severe acute respiratory syndrome-related coronavirus (SARS-CoV) epidemic (November 2002 – July 2003, Southern China), Middle East respiratory syndrome-related coronavirus (MERS-CoV) outbreaks (January 2014 – May 2014, Saudi Arabia and May 2015 – June 2015, South Korea), and the most recent Severe acute respiratory syndrome coronavirus 2 (SARS-CoV-2) worldwide pandemic (starting in November 2019 in Wuhan, China), this viral group is now extensively studied (Hung, 2003; Cowling et al., 2015; Oboho et al., 2015; Li et al., 2020).

The small size of RNA virus genomes is in principle linked to their limited ability for RNA synthesis, which is directly connected to a replication complex containing RNA-dependent RNA polymerase (RdRp) without reparation mechanisms (Gorbalenya et al., 2006). These RNA viruses can generate one mutation per genome per replication round (Drake and Holland, 1999). This combination of features means that RNA viruses are able to adapt to new environmental conditions, but they are limited in expanding their genomes because they must keep their mutation load low so that their survival is possible (Eigen and Schuster, 1977; Lauring et al., 2013; Carrasco-Hernandez et al., 2017). *Nidovirales* bind to their host cell receptors on the cell surface, after which fusion of the viral and cellular membranes is mediated by one of the surface glycoproteins. This mechanism, which progresses in the cytoplasm or endosomal membrane, releases the nucleocapsid into the host cell’s cytoplasm. After genome “transportation,” translation of two replicase open reading frames (ORFs) is initiated by host ribosomes. This results in large polyprotein precursors that undergo autoproteolysis to eventually produce a membrane-bound replicase/transcriptase complex. This complex initiates synthesis of the genome RNA and controls the synthesis of structural and some other proteins. New virus particles are assembled by association of the new genomes with the cytoplasmic nucleocapsid protein and subsequent envelopment of the nucleocapsid structure. Subsequently, the viral envelope proteins are inserted into intracellular membranes and targeted to the site of virus assembly (most often membranes between the endoplasmic reticulum and Golgi complex) and then they meet up with the nucleocapsid and trigger the budding of virus particles into the lumen of the membrane compartment. The newly formed virions then leave the cell by following the exocytic pathway toward the plasma membrane (Lai and Cavanagh, 1997; Gorbalenya, 2001; Snijder et al., 2003; Ziebuhr, 2004; Gorbalenya et al., 2006).

Today, SARS-CoV-2 is being studied intensively by scientists all over the world due to the ongoing 2020 coronavirus pandemic (Cohen, 2020). The origin of this virus is unknown, but recently a two-hit scenario was proposed wherein SARS-CoV-2 ancestors in bats first acquired genetic characteristics of SARS-CoV by incorporating a SARS-like receptor-binding domain through recombination prior to 2009; subsequently, the lineage that led to SARS-CoV-2 accumulated further, unique changes specifically within this domain (Patino-Galindo et al., 2020). As true of SARS-CoV, cell entry by SARS-CoV-2 depends upon angiotensin-converting enzyme 2 (ACE2) and transmembrane protease, serine 2 (TMPRSS2) for viral spike protein priming (Hoffmann et al., 2020). The genome of SARS-CoV-2 was sequenced and annotated in early January 2020 (Wu et al., 2020). A recent study revealed that the transcriptome of SARS-CoV-2 is highly complex due to numerous canonical and noncanonical recombination events. Moreover, it was found that SARS-CoV-2 produces transcripts encoding unknown ORFs and at least 41 potential RNA modification sites with an AAGAA motif were discovered in its RNA (Kim et al., 2020).

G-quadruplex binding proteins (QBPs) play crucial roles in many signaling pathways, including such biologically highly relevant activities as cell division, dysregulations of which lead to cancer development (Wu et al., 2008; Brázda et al., 2014). QBPs have been found to be involved in various viral infection pathways. An interesting example is HIV-1 nucleocapsid protein NCp7. It has been described how this protein helps to resolve an otherwise very stable G-quadruplex structure in viral RNA that stalls reverse transcription (Butovskaya et al., 2019). Various G-quadruplex-forming aptamers are used as drugs against many different viral proteins, suggesting a prominent role for QBP-mediated regulation. Among many other viruses, in particular Hepatitis C virus, HIV-1, and SARS-CoV are targets of these G-quadruplex-forming aptamers (Platella et al., 2017). Quadruplex binding domain has been found in nonstructural protein 3 (Nsp3) (Lei et al., 2018). This so-called SARS-UNIQUE-DOMAIN (SUD), and especially its M subdomain, was observed to be essential for SARS replication in host cells. Deletion or even substitution mutations in key RNA-interacting amino acids were shown to result in viral inability to replicate within host cells (Kusov et al., 2015). Moreover, this subdomain was found also in MERS and several other coronaviruses (Kusov et al., 2015).

G-quadruplexes are secondary nucleic acid structures formed in guanine-rich strands (Burge et al., 2006; Vorlíčková et al., 2012; Kolesnikova and Curtis, 2019). These have been detected in various genomes, but most extensively they have been described in human genomes (Chambers et al., 2015; Bedrat et al., 2016; Hänsel-Hertsch et al., 2016). They are present also in viruses (Lavezzo et al., 2018; Frasson et al., 2019). G-quadruplexes probably play an important role in regulating replication in most viral nucleic acids (Lavezzo et al., 2018), and these structures have been suggested as targets for antiviral therapy (Métifiot et al., 2014; Ruggiero and Richter, 2018). Along with cruciforms and hairpins, which can be formed in nucleic acids by inverted repeats (IRs), G-quadruplexes are significant genome elements playing specific biological roles. They are involved, for example, in the regulation of DNA replication and transcription (Bagga et al., 1990; Limanskaya, 2009; Brázda et al., 2011). It has been demonstrated that IRs are important for various processes in viruses, including their genome organization (Li and Li, 2010; Ishimaru et al., 2013; Xie et al., 2017; Bridges et al., 2019). Another interesting RNA motif that has been used as a drug target and was found in SARS-CoV targeted by 1,4-diazepame is the “slippery sequence” followed by a pseudoknot (Plant et al., 2005). This structure, common among all coronaviruses, works based on ribosomal −1 frameshifting that switches on viral fusion proteins (Plant et al., 2005).

In all prokaryotic and eukaryotic cells, as well as viruses, there have been found sequence motifs such as IR sequences forming cruciforms and hairpins or G-rich sequences that form G-quadruplexes (Brázda et al., 2011; Lavezzo et al., 2018; Bartas et al., 2019). In the present research, we conducted a systematic and comprehensive bioinformatic study searching for the occurrence of IRs and putative quadruplex sites (PQSs) within the genomes of all known *Nidovirales*. The aim was to find one or more potential druggable RNA targets to address the present COVID-19 threat (**Figure 1**).

**FIGURE 1.**
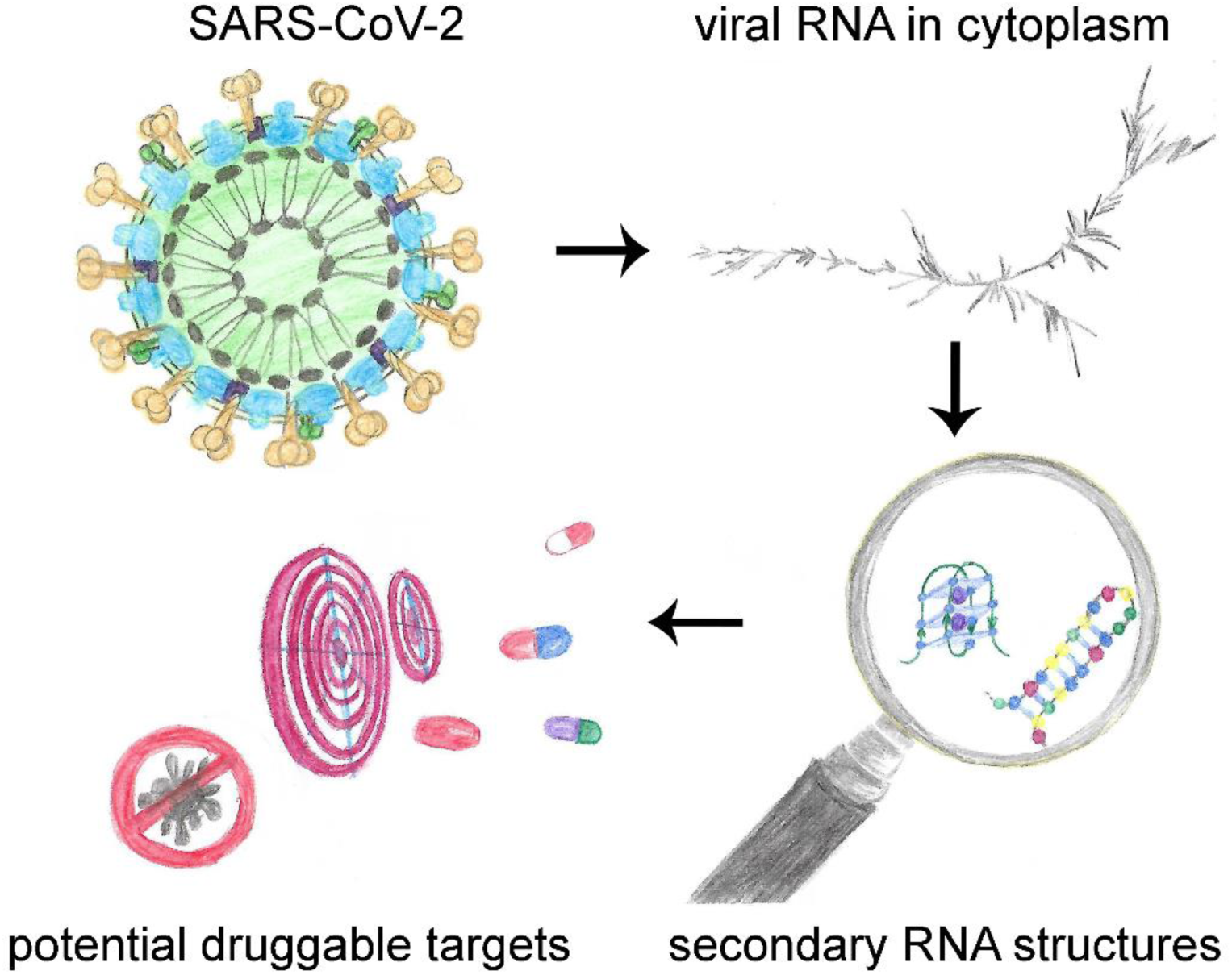
When outside host cells, the RNA of SARS-CoV-2 is highly spiralized by means of nucleocapsid phosphoproteins (top left). Once RNA is released into the cytoplasm and during replication/transcription processes, the formation of G-quadruplex and/or hairpins may take place (right). Stabilization of these structures by existing drugs may play a crucial role in inhibiting the viral lifecycle (bottom left).

## 2 Materials and Methods

### 2.1 Genome Source

Linear genomes of 109 viruses from the order *Nidovirales* were downloaded from the genome database of the National Center for Biotechnology Information (NCBI). Full names, phylogenetic groups, exact NCBI accession numbers, and further information are summarized in Supplementary Material 1.

### 2.2 Nidovirales Phylogenetic Tree Construction

Exact taxonomic identifiers of all analyzed *Nidovirales* species (obtained from Taxonomy Browser via NCBI Taxonomy Database [Drake and Holland, 1999]) were downloaded to phyloT: a tree generator (http://phylot.biobyte.de) and a phylogenetic tree was constructed using the function “Visualize in iTOL” in the Interactive Tree of Life environment (Letunic and Bork, 2019).

### 2.3 Analyses of PQSs Occurrence in *Nidovirales* Sequences

*Nidovirales* sequences were analyzed using our G4Hunter Web Tool (Brázda et al., 2019). The software is capable to read the NCBI identifier of the sequences uploaded in a .csv file. The parameters for G4Hunter were set to 25 as window size and G4Hunter score above 1.2. The results for each analyzed sequence contained information about the size of the genome and number of putative PQSs. All the results were merged into a single Microsoft Excel file where statistical analysis was then made. We also downloaded features tables of each virus from the NCBI database. These tables contain information about known features in the genome of each species. We searched the occurrence of G-quadruplex-forming sequences before, inside, and after the specific features of each genome using a predefined feature neighborhood ± 100 nucleotides (nt). Data were then plotted in Excel, where the statistical analysis was made. Complete analyses of PQSs occurrence in *Nidovirales* are provided in Supplementary Material 2.

### 2.4 Analyses of IRs Occurrence in *Nidovirales* Sequences

All *Nidovirales* genomes were analyzed by the core of our Palindrome analyser webserver (Brázda et al., 2016). The software was modified to read NCBI identifiers of sequences from text files. The size of IRs was set to 6–30 nt, size of spacers to 1–10 nt, and maximally one mismatch was allowed. A separate list of IRs in each of the 109 sequences and an overall report were exported. The overall report contained the lengths of the analyzed sequences, total number of IRs found, numbers of IRs grouped by size of IR (6–30 nt), and sum of IRs longer than 8 nt, 10 nt and 12 nt. The software also counted frequency of IRs in each sequence. Frequencies of IRs were normalized per 1,000 nt. Features tables of 109 *Nidovirales* genomes were downloaded from the NCBI database and grouped by their names as stated in the feature table file. Analyses of IRs occurrence inside and around (before and after) these features was performed. The search for IRs took place in predefined feature neighborhoods ± 100 nt around and inside feature boundaries. We calculated the numbers of all IRs and of those longer than 8, 10, and 12 nt in regions before, inside, and after the features. The categorization of an IR according to its overlap with a feature or feature neighborhood is demonstrated by the example shown in Supplementary Material 3. Complete analyses of IRs occurrence in *Nidovirales* are provided in Supplementary Material 4.

### 2.5 RNA Fold Predictions

In order to be able to display higher structures of the coronavirus genome, we used Galaxy’s free-online webserver (Afgan et al., 2018) and its RNA fold tool (Lorenz et al., 2011). This tool allows quick calculation of minimum free energy of secondary structures. We left the default parameters (Temperature 37 °C, Unpaired bases to participate in all dangling ends, Naview layout). We used SARS-CoV-2 genomic RNA sequence (NC_045512.2) in FASTA format as the input format. The output data were then displayed using the RNA plot tool [3]. We again left the default parameters (Naview layout, Output format Postscript .ps). We then downloaded the displayed secondary structures in high-resolution format. The raw data are provided in Supplementary Material 5.

### 2.6 Multiple Alignment of SUD Domains (M Regions) in Nsp3 of Pathogenic Species

Multiple protein alignment was done using MUSCLE (Edgar, 2004) under default parameters (UGENE [Okonechnikov et al., 2012] workflow was used). The following accessions were used: NP_828862.2 (Nsp3 SARS-CoV), YP_009047231.1 (Nsp3 MERS-CoV), and YP_009725299.1 (Nsp3 SARS-CoV-2). Conserved regions were added according to a graph published previously (Kusov et al., 2015).

### 2.7 Prediction of Human RNA-Binding Proteins Sites in SARS-CoV-2 RNA

The human SARS-CoV-2 RNA sequence was downloaded from NCBI (accession NC_045512.2) in FASTA format and inserted into the RBPmap (Mapping Binding Sites of RNA-binding proteins) web-based tool (Paz et al., 2014). The database of 114 human experimentally validated motifs was used. Both the “High stringency” and “Apply conservation filter” options were used. The output was further filtered in Excel to keep only those hits below *p*-value = 1.10^−6^. The complete results are provided in Supplementary Material 6.

## 3 Results

### 3.1 Phylogenetic Relationships in *Nidovirales*

According to the current state of knowledge, the order *Nidovirales* can be divided into six distinct families, and there are two unclassified species (**Figure 2**). *Coronaviridae* is the largest family and consists of 56 species. Among these are 12 species able to infect humans, including SARS-CoV, MERS-CoV, and SARS-CoV-2. Second largest is the *Arteriviridae* family with 22 species, third largest is the *Mesnidovirineae* family with 13 species, fourth is the *Tobaniviridae* family with 11 species, including 1 species able to infect humans. Fifth largest is the *Ronidovirineae* family with 5 species, and sixth is the *Mononiviridae* family with only 1 species.

**FIGURE 2.**
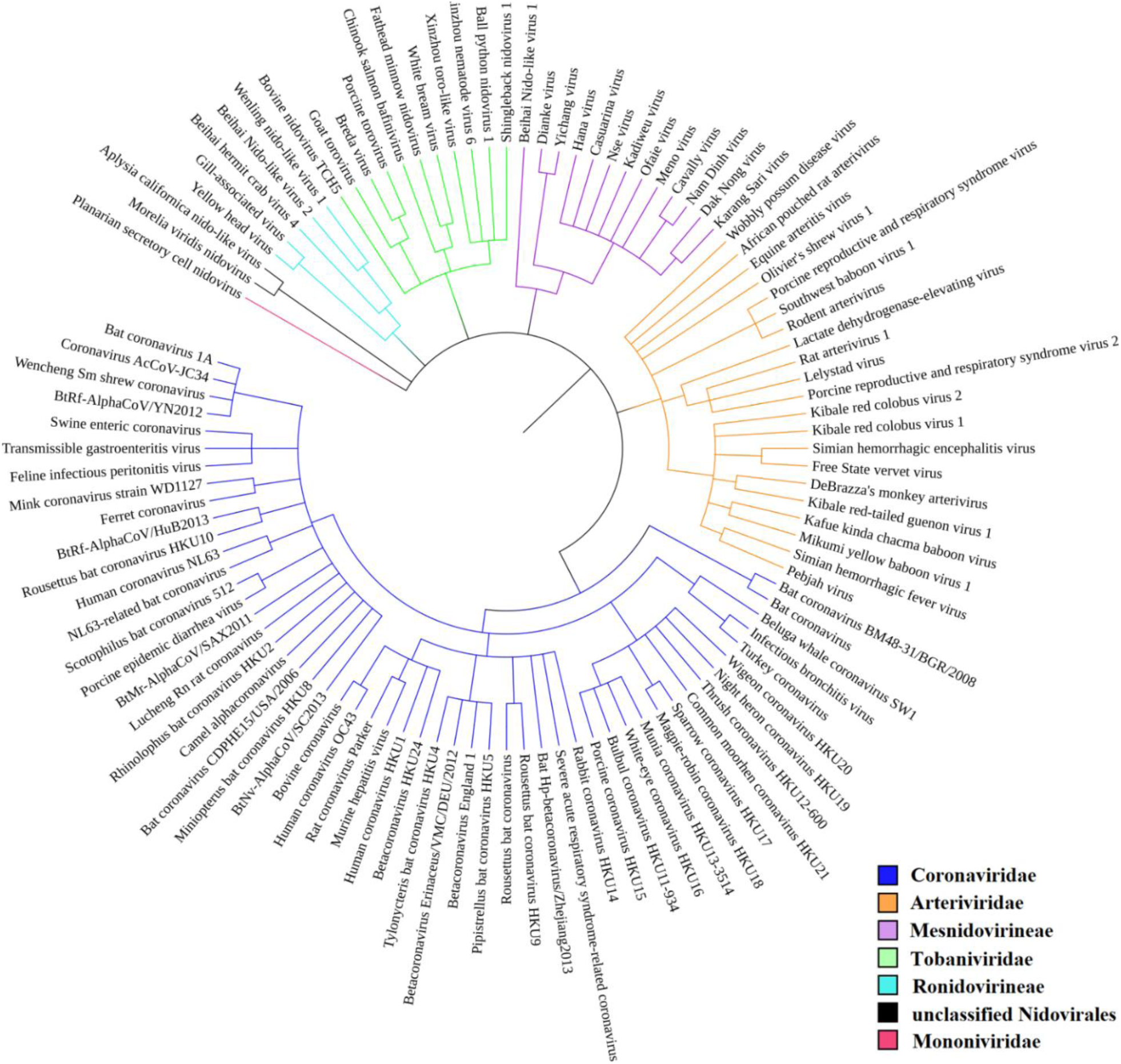
Phylogenetic relationships in *Nidovirales*. SARS-CoV-2 is not yet recognized by iToL, but, based upon current evidence, it probably will be placed close to the Bat Hp-betacoronavirus/Zheijang2013 and Severe acute respiratory syndrome-related coronavirus (SARS-nCoV) branches (Lu et al., 2020).

### 3.2 Variation in PQS Frequency in *Nidovirales*

We analyzed the occurrence of PQSs using G4Hunter in all 109 known genomes of *Nidovirales*. The length of genomes in the dataset varies from 12,704 nt (Equine arteritis virus) to 41,178 nt (Planarian secretory cell nidovirus). The mean GC content is 42.15%, the minimum is 27.50% for Planarian secretory cell nidovirus (*Mononiviridae* family), and the maximum is 57.35% for Beihai Nido-like virus 1 (*Mesnidovirineae* family). Using standard values for the G4Hunter algorithm (i.e., window size 25 and G4Hunter score above 1.2), we found 1,021 PQSs among all 109 *Nidovirales* genomes. The most abundant PQSs are those with G4Hunter scores of 1.2–1.4 (98.24% of all PQS), much less abundant are PQSs with G4Hunter scores 1.4–1.6 (1.76% of all PQS), and there were no PQSs above the 1.6 G4Hunter score threshold. In general, a higher G4Hunter score means a higher probability of G-quadruplexes forming inside the PQS (Bedrat et al., 2016). Genomic sequence sizes, total PQS counts, and PQS frequencies characteristics are summarized in **Table 1**.

**TABLE 1.** Genomic sequence sizes, PQS frequencies, and total counts. Seq (total number of sequences), Median (median length of sequences), Short (shortest sequence), Long (longest sequence), PQS (total number of predicted PQSs), Mean f (mean frequency of predicted PQS per 1,000 nt), Min f (lowest frequency of predicted PQS per 1,000 nt), Max f (highest frequency of predicted PQS per 1,000 nt), GC% (mean GC content). Note that 132 PQSs were in *Nidovirales* with Undefined host.

| All | Seq | Median | Short | Long | PQS | Mean f | Min f | Max f | GC% |
| --- | --- | --- | --- | --- | --- | --- | --- | --- | --- |
| Nidovirales | 109 | 26,975 | 12,704 | 41,178 | 1,021 | 0.48 | 0 | 9.07 | 42.15 |
| Family | Seq | Median | Short | Long | PQS | Mean f | Min f | Max f | GC% |
| Arteriviridae | 22 | 15,236 | 12,704 | 15,728 | 392 | 1.18 | 0.45 | 2.72 | 51.60 |
| Coronaviridae | 56 | 28,345 | 20,398 | 31,686 | 162 | 0.10 | 0 | 0.52 | 39.39 |
| Mesnidovirineae | 12 | 20,117 | 19,867 | 20,949 | 250 | 1.03 | 0 | 9.07 | 38.75 |
| Ronidovirineae | 5 | 26,253 | 24,648 | 29,385 | 51 | 0.39 | 0.04 | 1.07 | 44.79 |
| Tobaniviridae | 11 | 27,318 | 20,261 | 33,452 | 129 | 0.43 | 0 | 1.48 | 40.94 |
| By hosts | Seq | Median | Short | Long | PQS | Mean f | Min f | Max f | GC% |
| Invertebrates | 17 | 20,307 | 19,917 | 41,178 | 298 | 0.82 | 0 | 9.07 | 39.76 |
| Vertebrates (including Humans) | 83 | 27,608 | 12,704 | 33,452 | 591 | 0.38 | 0 | 1.75 | 42.74 |
| Humans | 13 | 29,751 | 27,317 | 31,028 | 21 | 0.06 | 0 | 0.32 | 37.88 |
| SARS-CoV-2 | 1 | 29,903 | - | - | 1 | 0.03 | - | - | 37.97 |

The highest PQS frequencies were found in Beihai Nido-like virus 1, which is an intracellular parasite of sea snails in genus *Turritella*. Beihai Nido-like virus 1 has in total 184 PQSs in its genomic sequence 20,278 nt long and PQS frequency of 9.07 PQS per 1,000 nt. On the other hand, no PQSs were found in the genomic sequences of 16 other *Nidovirales* species. The mean PQS value for all *Nidovirales* was 0.48 PQS per 1,000 nt. By viral families, the highest mean PQS frequency per 1,000 nt was in *Arteriviridae* (1.18), followed by in *Mesnidovirineae* (1.03), much lower in *Tobaniviridae* (0.43) and in *Ronidovirineae* (0.39), and the lowest was in *Coronaviridae* (0.10). When grouped and analyzed by host organisms, new information became apparent. The highest mean PQS frequency was in *Nidovirales* infecting Invertebrates (0.80), then in *Nidovirales* infecting Vertebrates including Humans (0.38), and the last in *Nidovirales* infecting Humans (0.06). In pathogenic human coronaviruses, the total PQSs counts were as follow: 4 PQS in SARS-CoV, 0 PQS in MERS-CoV, and 1 PQS in SARS-CoV-2. To test whether the PQS frequency per 1,000 nt in SARS-CoV-2 is significantly greater than in a random sequence, we generated 10 random sequences with length and GC content the same as in SARS-CoV-2. In that test, the mean PQS frequency per 1,000 nt was 0.16 with standard deviation of 0.05. We applied Wilcoxon signed-rank test with continuity correction and the resulting *p*-value was 0.0028. Thus, the PQS frequency per 1,000 nt in SARS-CoV-2 (0.03) is significantly lower than expected.

All PQSs found in ranges of G4Hunter score intervals and precomputed PQS frequencies per 1,000 nt are summarized in **Table 2**. The relationship between observed PQS frequency per 1,000 nucleotides and GC content in all analyzed *Nidovirales* sequences is depicted in **Figure 3**, the Beihai Nido-like virus has both the highest GC content and highest PQS frequency. Within *Nidovirales* with Humans as host, however, the highest PQS frequency was found in Breda virus, which does not have the highest GC content in the dataset.

**TABLE 2.** Total number of PQSs and their resulting frequencies per 1,000 nt in all 109 genomes of *Nidovirales* and in particular categories according to their hosts, grouped by G4Hunter score. Frequency was computed using total number of PQSs in each category divided by total length of all analyzed sequences and multiplied by 1,000. Note that 132 PQSs were in *Nidovirales* with Undefined host.

| Interval of G4Hunter Score | Number of PQSs in Dataset | PQS Frequency per 1,000 nt |
| --- | --- | --- |
| **All** |  |  |
| 1.2 | 1,003 | 0.47 |
| 1.4 | 18 | 0.01 |
| **Arteriviridae** |  |  |
| 1.2 | 386 | 1.17 |
| 1.4 | 6 | 0.02 |
| **Coronaviridae** |  |  |
| 1.2 | 157 | 0.10 |
| 1.4 | 5 | 0.003 |

| Mesnidovirineae |  |  |
| --- | --- | --- |
| 1.2 | 245 | 1.01 |
| 1.4 | 5 | 0.02 |
| Ronidovirineae |  |  |
| 1.2 | 50 | 0.38 |
| 1.4 | 1 | 0.01 |
| Tobaniviridae |  |  |
| 1.2 | 129 | 0.43 |
| 1.4 | 0 | 0 |
| Humans only |  |  |
| 1.2 | 20 | 0.05 |
| 1.4 | 1 | 0.003 |
| Vertebrate<br>(Including Humans) |  |  |
| 1.2 | 580 | 0.37 |
| 1.4 | 11 | 0.01 |
| Invertebrates |  |  |
| 1.2 | 292 | 0.80 |
| 1.4 | 6 | 0.02 |

**FIGURE 3.**
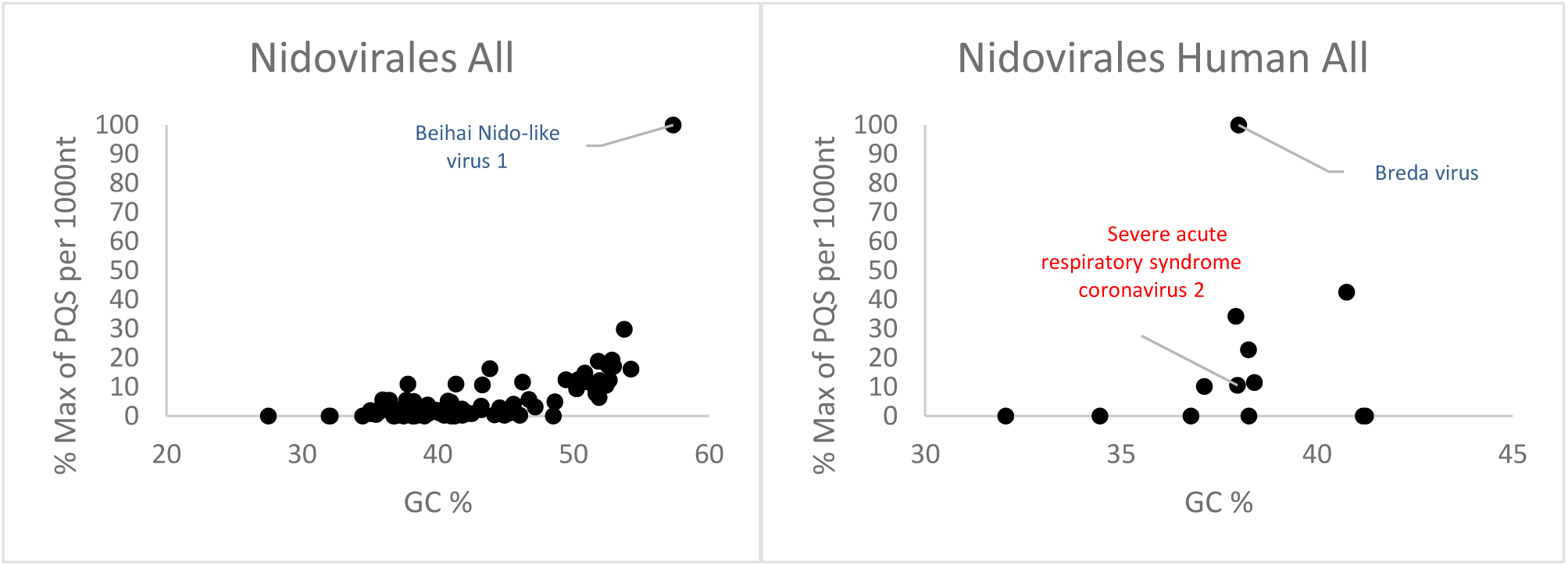
Relationship between observed PQS frequency per 1,000 nucleotides and GC content in all analyzed *Nidovirales* sequences. In each G4Hunter score interval miniplot, frequencies were normalized according to the highest observed PQS frequency. Organisms with maximum frequency per 1,000 nt greater than 50% are described and highlighted in color

Most of the PQSs have G4Hunter scores between 1.2 and 1.4. Only a few sequences have G4Hunter scores above 1.4. In comparison with other *Nidovirales*, the members of the *Coronaviridae* and especially pathogenic human *Coronaviridae* have the lowest PQS frequency in the dataset.

**Figure 4** shows the differences in PQS frequency by annotated loci. We downloaded features for every virus genome and analyzed the presence of PQS in each annotated sequence and in its proximity (before and after). The most notable enrichment of PQSs was located before 3′UTR, inside 5′UTR, before mRNA, and after miscellaneous RNA. The lowest PQS frequencies were found after 3′UTR, before 5′UTR, before and inside miscellaneous RNA, and in miscellaneous features.

**FIGURE 4.**
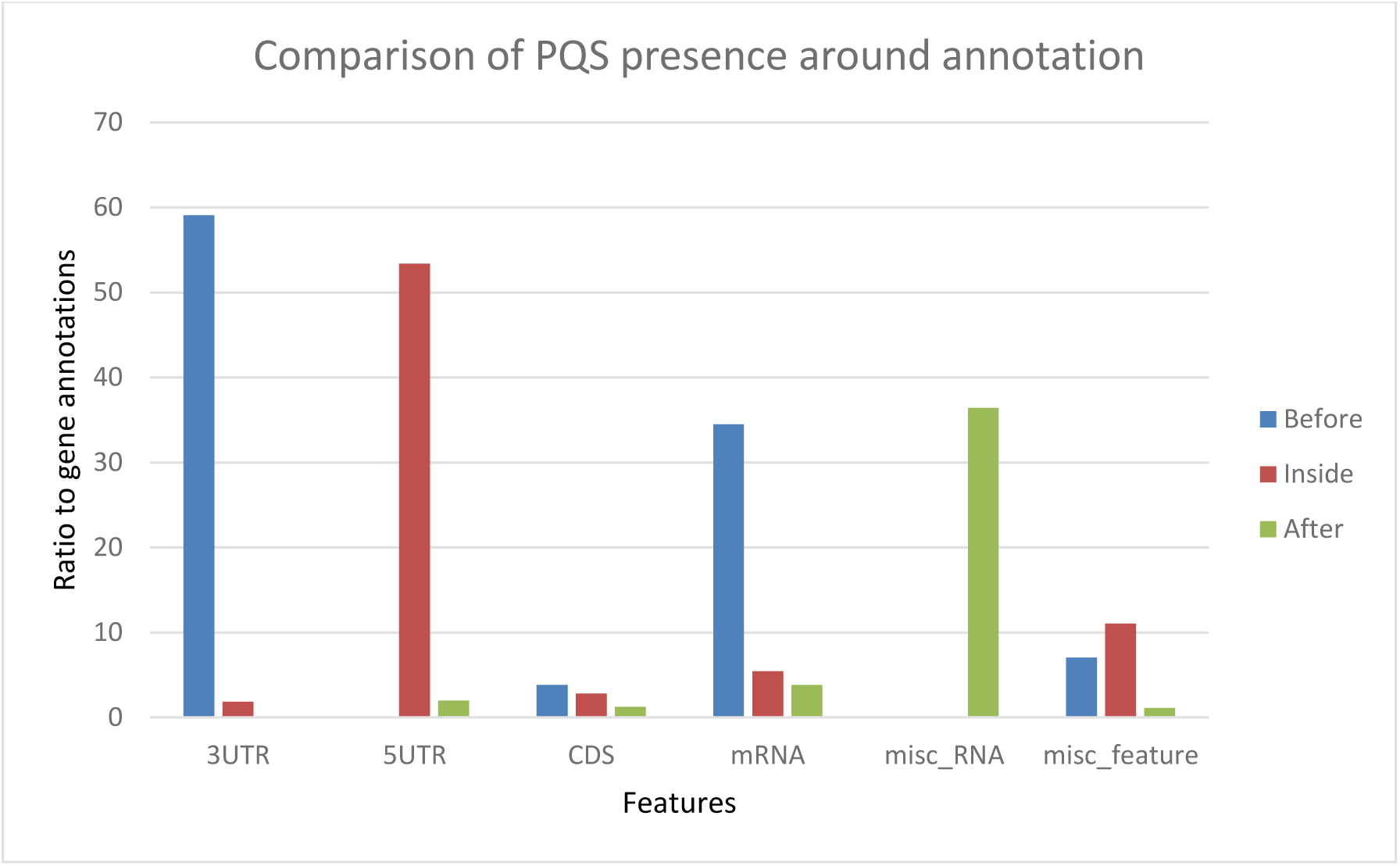
Differences in PQS frequency by annotated locus. The figure shows the PQS frequencies between annotations from the NCBI database. We analyzed frequencies of all PQSs inside, before, and after the annotations.

### 3.3 Variation in Frequency of Inverted Repeats in *Nidovirales*

To find short IRs in *Nidovirales* genomic sequences, we utilized the core of the *Palindrome analyser* (Brázda et al., 2016). We used values for *Palindrome analyser* returning inverted repeats capable to form stable hairpins (size of IRs was set to 6–30 nt, size of spacer to 1–10 nt, maximally one mismatch). Genomic sequence sizes, total IR counts, and IR frequencies characteristics are summarized in **Table 3**. In total, 93,369 IRs were found and the mean IRs frequency was 33.90 per 1,000 nt. The maximal IR frequency was found in Planarian secretory cell nidovirus (56.17), the species with the largest genome among *Nidovirales*. It has been suggested that Planarian secretory cell nidovirus diverged early from multi-ORF *Nidovirales* and acquired additional genes, including those typical of large DNA viruses or hosts (RNAse T2, Ankyrin, and Fibronectin type II), which might modulate virus–host interactions (Saberi et al., 2018). The lowest IR frequency was noticed in Beihai Nido-like virus 1 (19.23). Noteworthy is that this is the species of *Nidovirales* with the highest GC content and PQS frequency. Differences in IR frequency by annotated locus are depicted in **Figure 5**. By families, the highest mean IRs frequency was found in *Coronaviridae* (37.08) and lowest in *Ronidovirineae* (26.37). Novel human coronavirus SARS-CoV-2 has relatively high IR frequency per 1,000 nt, in comparison with other *Nidovirales*. To test if the frequency of IRs per 1,000 nt in SARS-CoV-2 is significantly greater than in random sequence, we generated 10 random sequences with length and GC content the same as in SARS-CoV-2. The mean frequency of IRs per 1,000 nt was 30.23, with standard deviation 0.22. We applied Wilcoxon signed-rank test with continuity correction to find that *p*-value was equal to 0.0029. Thus, the IR frequency per 1,000 nt in SARS-CoV-2 (40.23) is significantly higher than expected. When we inspected differences of IR frequencies according to hosts, we found the highest mean IR frequency per 1,000 nt to be in *Nidovirales* infecting Humans (38.13), then in Vertebrates (34.25), and the lowest mean frequency is in Invertebrates (31.77).

**TABLE 3.** Genomic sequence sizes, IRs frequencies and total counts. Seq (total number of sequences), Median (median length of sequences), Short (shortest sequence), Long (longest sequence), GC% (mean GC content), IRs (total number of predicted IRs), Mean f (mean frequency of predicted IRs per 1,000 nt), Min f (lowest frequency of predicted IRs per 1,000 nt), and Max f (highest frequency of predicted PQS per 1,000 nt) are shown for all 109 *Nidovirales* sequences, families, host groups, and for SARS-CoV-2 alone.

| Order | Seq | Median | Short | Long | IRs | Mean f | Min f | Max f | GC% |
| --- | --- | --- | --- | --- | --- | --- | --- | --- | --- |
| Nidovirales | 109 | 26,975 | 12,704 | 41,178 | 93,369 | 33.90 | 19.23 | 56.17 | 42.15 |
| Family | Seq | Median | Short | Long | IRs | Mean f | Min f | Max f | GC% |
| Arteriviridae | 22 | 15,236 | 12,704 | 15,728 | 9,695 | 29.44 | 26.47 | 33.57 | 51.60 |
| Coronaviridae | 56 | 28,345 | 20,398 | 31,686 | 58,943 | 37.08 | 31.13 | 48.16 | 39.39 |
| Mesnidovirineae | 12 | 20,117 | 19,867 | 20,949 | 8,012 | 33.04 | 19.23 | 39.06 | 38.75 |
| Ronidovirineae | 5 | 26,253 | 24,648 | 29,385 | 3,474 | 26.37 | 22.20 | 29.62 | 44.79 |
| Tobaniviridae | 11 | 27,318 | 20,261 | 33,452 | 9,082 | 30.24 | 23.96 | 37.95 | 40.94 |
| Host | Seq | Median | Short | Long | IRs | Mean f | Min f | Max f | GC% |
| Invertebrates | 17 | 20,307 | 19,917 | 4,178 | 13,233 | 31.77 | 19.23 | 56.17 | 39.76 |
| Vertebrates<br>(including Humans) | 83 | 27,608 | 12,704 | 33,452 | 45,844 | 34.25 | 24.18 | 48.16 | 42.74 |
| Humans | 13 | 29,751 | 27,317 | 31,028 | 14,408 | 38.13 | 33.89 | 43.44 | 37.88 |
| SARS-CoV-2 | 1 | 29,903 | - | - | 1,203 | 40.23 | - | - | 37.97 |

**FIGURE 5.**
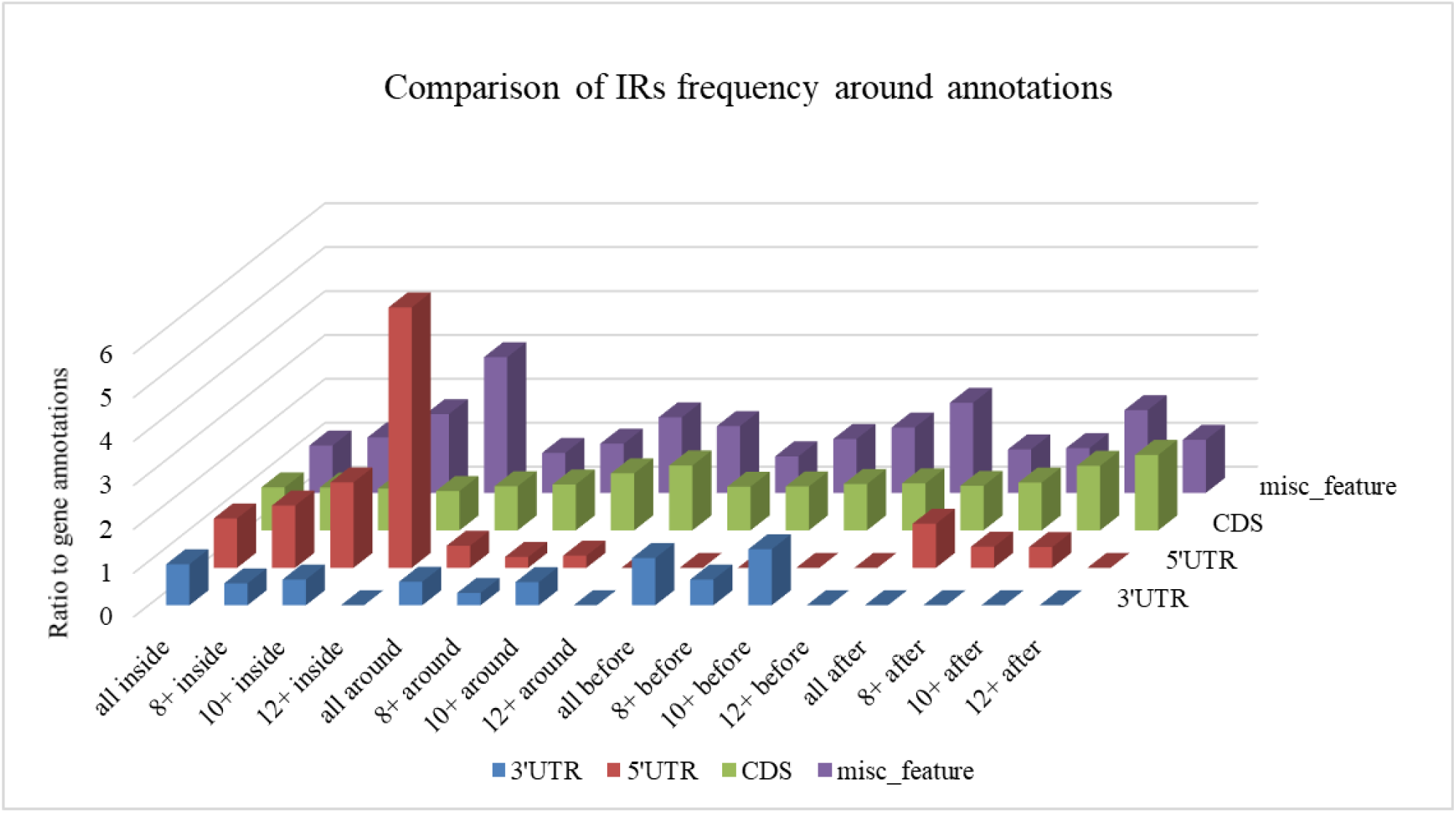
Differences in IR frequency by annotated locus. The chart compares IR frequencies per 1,000 nt between “gene” annotation and other annotated locations from the NCBI database that were found in genomes more than 10 times. We analyzed frequencies of all IRs (all) and of IRs with lengths 8 nt and longer (8+), 10 nt and longer (10+,) and 12 nt and longer (12+) within annotated locations (inside) as well as before and after annotated locations.

A summary of all IRs found in ranges of different IR sizes and precomputed IR frequencies per 1,000 nt is shown in **Table 4**. Although generally the frequency of IR presence decreases with the IR length, there are notable differences between groups and also between viruses with different hosts. The most as well as longest IRs occur in the *Coronaviridae* group and in viruses having humans as a host. IRs 12 bp long and longer are very rare in the *Ronidovirineae* group. The relationship between IRs length and stability of resulting secondary structure is not simple. While some authors believe that longer IRs are more stable, others suggest that there is an energy optimum defined by arm and spacer length (Sinden et al., 1991; Brázda et al., 2016; Georgakopoulos-Soares et al., 2018).

**TABLE 4.** Total number of IRs and their resulting frequencies per 1,000 nt grouped by size of IR. Shown are data for all 109 representative *Nidovirales* sequences, for families, host groups, and for SARS-CoV-2 alone. Frequency was computed using total number of IRs in each category divided by total length of all analyzed sequences and multiplied by 1,000.

| Size of IRs | Number of IRs in Dataset | IRs Frequency per 1,000 nt |
| --- | --- | --- |
| **All** |  |  |
| All | 93,369 | 33.90 |
| 8+ | 12,669 | 4.54 |
| 10+ | 1,872 | 0.65 |
| 12+ | 244 | 0.08 |
| **Arteriviridae** |  |  |
| All | 9,695 | 29.44 |
| 8+ | 1,127 | 3.41 |
| 10+ | 103 | 0.31 |
| 12+ | 8 | 0.02 |
| **Coronaviridae** |  |  |
| All | 58,943 | 37.08 |
| 8+ | 8,527 | 5.36 |
| 10+ | 1,393 | 0.88 |
| 12+ | 190 | 0.12 |
| **Mesnidovirineae** |  |  |
| All | 8,012 | 33.04 |
| 8+ | 1,081 | 4.46 |
| 10+ | 137 | 0.56 |
| 12+ | 13 | 0.05 |
| **Ronidovirineae** |  |  |
| All | 3,474 | 26.37 |
| 8+ | 320 | 2.44 |
| 10+ | 25 | 0.18 |
| 12+ | 2 | 0.01 |
| **Tobaniviridae** |  |  |
| All | 9,082 | 30.24 |
| 8+ | 1,060 | 3.54 |
| 10+ | 128 | 0.43 |
| 12+ | 15 | 0.05 |
| **Humans only** |  |  |
| All | 14,408 | 38.13 |
| 8+ | 2,048 | 5.40 |
| 10+ | 344 | 0.91 |
| 12+ | 42 | 0.11 |
| **Vertebrate**<br>**(Including Humans)** |  |  |
| All | 72,838 | 34.25 |
| 8+ | 10,097 | 4.66 |
| 10+ | 1,549 | 0.69 |
| 12+ | 207 | 0.09 |
| **Invertebrates** |  |  |
| All | 13,233 | 31.77 |
| 8+ | 1,686 | 4.06 |
| 10+ | 218 | 0.51 |
| 12+ | 24 | 0.05 |
| **SARS-CoV-2** |  |  |
| All | 1,203 | 40.23 |
| 8+ | 203 | 6.79 |
| 10+ | 37 | 1.24 |
| 12+ | 5 | 0.17 |

Differences in IR frequency according to annotated loci are shown in **Figure 5**. The most notable enrichment of IRs was found inside 5′UTR for IRs of size 12+ nt, and this is the most frequently occurring location of 12+ IRs in *Nidovirales* genomes. Noteworthy is that there are no 12+ IRs around 5′UTR loci, but there is an abundance of IRs 10+ nt long in these locations. The 5′UTR are abundant for 12+ IR, but there are no 12+ IRs within 3′UTR. This points to functional relevance of these IRs in viral genomes.

### 3.4 Prediction of Human SARS-CoV-2 RNA Folding

It is very likely that coronavirus RNA in the viral capsid is compactly folded into a spiral-like structure due to nucleocapsid phosphoprotein N, as was proposed earlier for SARS-CoV (Chang et al., 2014). But what structure does the RNA takes after the capsid content is released into the cytoplasm of the host cell? We made a global prediction of RNA folding for SARS-CoV-2 and in a negative control (random RNA sequences with the same GC content and length). It was clearly visible that there arose a much more compact RNA structure in SARS-CoV-2 (**Figure 6**).

**FIGURE 6.**
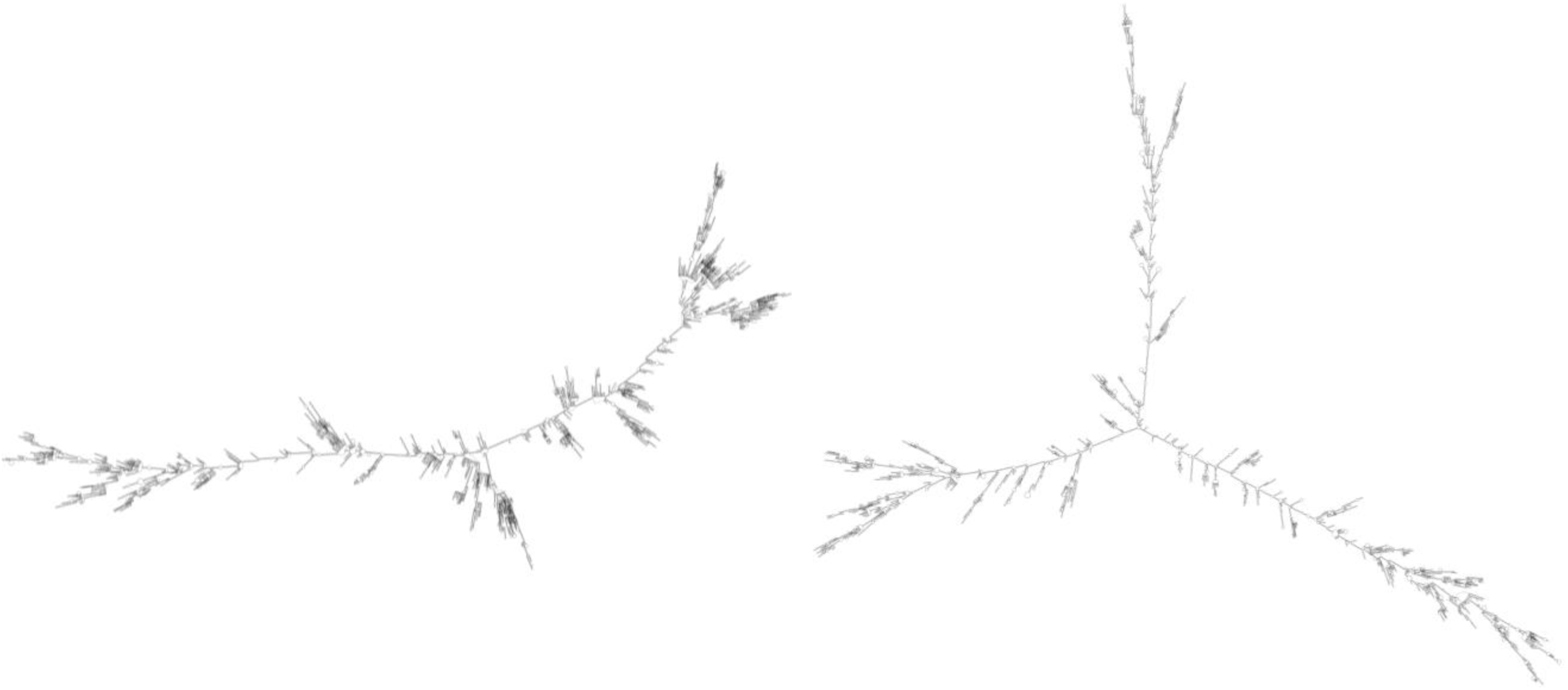
RNA fold (Lorenz et al., 2011) prediction for SARS-CoV-2 RNA molecule (left) and random sequence negative control (right). This figure shows a high level of SARS-CoV-2 genome folding via complementarity of particular RNA regions and forming of hairpins and/or cruciforms. RNA fold prediction was carried out using default parameters via Galaxy webserver (Afgan et al., 2018), which enables queries of lengths longer than 10,000 nucleotides.

**FIGURE 7.**
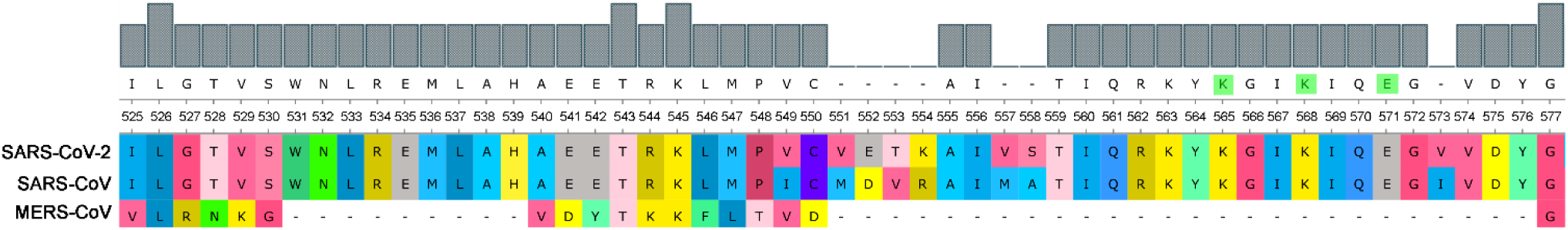
Comparison of SUD domain M region in three pathogenic human coronaviruses. Nsp3 proteins of three pathogenic human viruses (SARS-CoV-2, SARS-CoV, and MERS-CoV) were aligned and part of the critical G-quadruplex binding M region of the SUD domain (525–577 aa according to the SARS-CoV reference sequence) was visualized. As predicted according to Kusov et al. (2015), the G-quadruplex-interacting residues K565, K568, and E571 are highlighted in green (upper consensus panel).

### 3.5 Comparison of SUD Domain M Region of Nsp3 in Three Pathogenic Human Coronaviruses

Consisting of nearly 2,000 amino acids, Nsp3 is the largest multi-domain protein in *Nidovirales.* Nevertheless, Nsp3’s role is still largely unknown. It is believed that the protein plays various roles in coronavirus infection. Nsp3 interacts with other viral Nsps as well as RNA to form the replication/transcription complex. It also acts on posttranslational modifications of host proteins to antagonize the host innate immune response. On the other hand, Nsp3 is itself modified in host cells by N-glycosylation and can interact with host proteins to support virus survival (Lei et al., 2018). Both SARS-CoVs (2003 and 2019) have the PQSs in their genomes and have a retained M region of the SUD domain in protein Nsp3 that is reported to be critical for interacting with G-quadruplexes (Kusov et al., 2015). MERS has no PQS in its RNA sequence and, strikingly, it has a deletion in this region of the SUD domain suggesting parallel evolution simplifications.

### 3.6 Prediction of Human RNA-Binding Proteins Sites in SARS-CoV-2 RNA

Using the RBPmap tool, we predicted human RNA-binding proteins sites in SARS-CoV-2 RNA. While holding to very stringent thresholds, three highly promising candidate human RNA-binding proteins were predicted. SRSF7 was predicted to bind viral RNA in nucleotide position 26,194–26,199 (NC_045512.2), which is in fact exactly the binding motif for this protein (ACGACG). Protein HNRNPA1 was predicted to bind viral RNA in nucleotide position 23,090–23 097 (exact binding motif GUAGUAGU). The last protein found was TRA2A in nucleotide position 3,056–3,065 and position 3,074–3,083, in both cases with one mismatch (the motifs found were GAAGAAGAAG and the experimentally validated motif for TRA2A is GAAGAGGAAG). All three of these proteins share multiple RGG-rich novel interesting quadruplex interaction (NIQI) motifs, which are common to most human G-quadruplex-binding proteins (Brázda et al., 2018; Huang et al., 2018; Masuzawa and Oyoshi, 2020). All find individual motif occurrence (FIMO) hits below *p*-value = 1.10^−4^ are enclosed in Supplementary Material 7. The proteins SRSF7, HNRNPA1, and TRA2A are involved in mRNA splicing via the spliceosome pathway (Szklarczyk et al., 2015).

## 4 Discussion

Local DNA structures have been shown to play important roles in basic biological processes, including replication and transcription (Brázda et al., 2011; Travers and Muskhelishvili, 2015; Surovtsev and Jacobs-Wagner, 2018). PQS analyses of human viruses have clearly demonstrated that these sequences are conserved also in the viral genomes (Lavezzo et al., 2018) and could be targets for antiviral therapies (Wang et al., 2015; Krafčíková et al., 2017; Ruggiero and Richter, 2018). For example, the conserved PQS sequence present in the L gene of Zaire ebolavirus and related to its replication is inhibited by interaction with G-quadruplex ligand TMPyP4, and this finding has led to suggestions that G-quadruplex RNA stabilization could constitute a new strategy against Ebola virus disease (Wang et al., 2016). Our results found only one PQS in the genomic sequence of SARS-CoV-2, that having an absolute G4Hunter score of 1.24 and the following sequences: 5′-CCCCAAAAUCAGCGAAAUGCACCCC-3′ for a positive-strand intermediate and 5′-GGGGTGCATTTCGCTGATTTTGGGG-3′ for a negative-strand intermediate (where G-quadruplex could theoretically arise). This PQS is located in the position 28,289–28,313 within the nucleocapsid phosphoprotein coding sequence. It is noteworthy that all viral RNAs are produced through negative-strand intermediates, which are only about 1% as abundant as their positive-sense counterparts (Fehr and Perlman, 2015). It would be of great value to know whether TMPyP4 or other G-quadruplex stabilizing compounds can inhibit replication processes of SARS-CoV-2. RNA hairpins, which are formed by IRs, are basic structural elements of RNA and play crucial roles in gene expression and intermolecular recognition. Conserved palindromic RNA structures have been found in many viral genomes, including HIV-1, and play a crucial role in their replication (Liu et al., 2018). We have found an abundance of IRs inside 5′UTR in *Nidovirales* genomes. In general, 5′UTR is an important locus for regulation of viral replication and gene expression. It has been demonstrated that stem integrity of phylogenetically conserved stem-loop structure located in 5′UTR of the PRRSV virus from the *Arteriviridae* family is crucial for viral replication and subgenomic mRNA synthesis. Similar secondary structures have been proposed for several viruses from the *Arteriviridae* and *Coronaviridae* families (Lu et al., 2011). Our analyses of annotated features in **Figure 5** therefore support this report. The discrete locations of specific IRs in viral genomes could therefore be additional targets for their regulation.

Targeting viral proteins is usually effective only against specific viral strains and fails even for closely related viral species. It appears that targeting host proteins should be able to provide a response toward a wider spectrum of viruses inasmuch as different viruses exploit common cellular pathways. Many cellular RNA-binding proteins (RBPs) containing well-established RNA-binding domains (RBDs) are known to be critical for infection by different viruses. Recently, 472 RBPs were reviewed for their linkage to viruses (Garcia-Moreno et al., 2018). It has been demonstrated that G-quadruplex formation in HIV-1 viral genome stalls RNA polymerase, thereby limiting viral replication in host cells. HIV-1 nucleocapsid protein NCp7 helps to resolve G-quadruplex formation and therefore enables virus to spread. Stabilization of this quadruplex has been targeted by several experimental compounds. This treatment slowed or inhibited viral growth (Butovskaya et al., 2019).

Virus reproduction is dependent on the cellular transcription machinery, and therefore the interaction of the cellular proteins with viral RNA could be another target for antiviral therapy (Roberts et al., 2009). Our analysis predicted several QBPs to be capable to bind SARS-CoV-2 RNA. One of these, HNRNPA1 protein, is responsible for nuclear–cytoplasm shuttling (Garcia-Moreno et al., 2018). Like some other RNA-binding proteins, HNRNPA1 forms so-called membraneless organelles. These organelles are assemblies of proteins along with RNA or DNA that condense in specific cellular loci. The organelles can undergo transition from liquid-like droplets to amyloid fibrils, and mutations in so-called low complexity domains of these proteins lead to formation of amyloid aggregations that are found in many neurodegenerative diseases (Gui et al., 2019). HNRNPA1 is involved in many different cellular processes and has been targeted in various diseases. One such example is the varicella zoster virus that causes chicken pox. Moreover, this response to varicella zoster virus has been connected with autoimmune disease that complicates multiple sclerosis (Kattimani and Veerappa, 2018). HNRNPA1 is known to bind and resolve G-quadruplex formed in TRA2B promoter and to promote its transcription. Dysregulation of this binding leads to progression of colon cancer. This interaction has been targeted by the well-known G-quadruplex stabilizer pyridostatin, which led to decreased transcription from the TRA2B promoter (Nishikawa et al., 2019). Furthermore, HNRNPA1 is co-expressed with bromodomain and extraterminal domain protein BRD4 in human tumor samples. It has been shown experimentally that the well-described, naturally occurring polyphenolic flavonoid quercetin inhibited this protein and thereby led to better susceptibility to treatment in cancer patients (Pham et al., 2019). Both SRSF7 and TRA2A proteins, which are predicted to interact with SARS-CoV-2 RNA, play roles in alternative splicing (Ghosh et al., 2016). SRSF7 is a serine and arginine-rich splicing factor and is part of the spliceosome. Its expression has been connected to several types of cancer and it has been shown that its knockdown induced p21 expression and thus reduced cancer development (Saijo et al., 2016). Little is known about TRA2A protein. It was first identified in insects together with its paralog TRA2B, which has been studied to a greater extent (Tan et al., 2018). It has been found that TRA2A can promote paclitaxel therapy and promote cancer progression in triple-negative breast cancers (Liu et al., 2017). A role of TRA2A protein in regulation of HIV1 virus replication has been described. Both TRA2A and TRA2B bind to a specific HIV1 sequence and regulate its replication within the cell through alternative splicing of viral RNA (Erkelenz et al., 2013).

Scientists around the world are united in their efforts to find an effective therapy against coronavirus disease (COVID-19). Among the most promising candidates are remdesivir and chloroquine. Remdesivir is an adenosine analogue that incorporates into nascent viral RNA chains and results in premature termination. Preliminary data have shown that remdesivir effectively inhibited virus infection in a human liver cancer cell line (Wang et al., 2020). Chloroquine is a potential broad-spectrum antiviral drug and it already has been widely used as a low-cost and safe anti-malarial and autoimmune disease drug for more than 70 years. Application of chloroquine causes elevation of endosomal pH and also interferes with terminal glycosylation of the cellular receptor ACE2. This probably has a negative influence on virus-receptor binding and abrogates the infection (Vincent et al., 2005). Drug repurposing seems like a very good strategy for quickly and at low cost finding a new therapy for new human diseases (Oprea and Mestres, 2012). To date, there are few substances with G-quadruplex stabilizing features. One example is topotecan (also known by its brand name Hycamtin®), which frequently is used for treating ovary cancer. If everything else fails, it seems to be an option. On the other hand, topotecan cause unpleasant side effects, such as nausea, vomiting, and diarrhea (Topotecan - Chemotherapy Drugs - Chemocare). Many studies have described strong G-quadruplex stabilization effects (Li et al., 2018; Satpathi et al., 2018), which might be one possible mode of action. Moreover, it has been proven that higher structures of nucleic acids, and G-quadruplex especially, might be stabilized by use of various natural substances. Berbamine is one such substance and is a component of traditional Chinese medicine. It frequently is used for treating chronic myeloid leukemia or melanoma and has strong binding affinity to G-quadruplex structures (Tan et al., 2014). Viral nucleic acids and their loci with G-quadruplex-forming potential are in all cases very specific and are promising molecular targets for treating serious diseases. Evidence of G-quadruplex formation as a potential target for therapy was proven for Hepatitis A, flu virus, and HIV-1 (Métifiot et al., 2014). Because all the aforementioned cases concern RNA viruses, the viral G-quadruplexes are localized in the cytoplasm of the host cell. One of the main conclusions to be drawn from this study is that sequences able to form G-quadruplex structures in the *Nidovirales* (infecting human) family are very rare, and this suggests intentional suppression during evolution to simplify viral RNA replication. Finding the proper stabilizers of viral higher RNA structures might be crucial for inhibiting or stopping viral RNA replication in order to gain time for the immune system to deal successfully with an infection.

## Supporting information

Supplementary Material 1: Summary of analyzed Nidovirales genomes (full names, phylogenetic groups, exact NCBI accession, and further information)

Supplementary Material 2: Complete analyses of PQS occurrence in Nidovirales

Supplementary Material 3: Categorization of IRs according to their overlap with a feature or feature neighborhood

Supplementary Material 4: Complete analyses of IRs occurrence in Nidovirales

Supplementary Material 5: RNA fold prediction for SARS-CoV2 RNA and random RNA of the same length and GC content

Supplementary Material 7: Prediction of RGG-rich NIQI motifs in proteins identified by RBPmap

## 7 Acknowledgments

We would like to express our gratitude to Alena Volná, M.Sc. for her time spent in preparing the illustrations accompanying this work.

## Supplementary Materials

**Supplementary Material 1**: Summary of analyzed *Nidovirales* genomes (full names, phylogenetic groups, exact NCBI accession, and further information)

**Supplementary Material 2:** Complete analyses of PQS occurrence in *Nidovirales*

**Supplementary Material 3:** Categorization of IRs according to their overlap with a feature or feature neighborhood

**Supplementary Material 4:** Complete analyses of IRs occurrence in *Nidovirales*

**Supplementary Material 5:** RNA fold prediction for SARS-CoV2 RNA and random RNA of the same length and GC content

**Supplementary Material 6:** Complete RBPmap results – Prediction of human RNA-binding protein sites in SARS-CoV-2 RNA

**Supplementary Material 7:** Prediction of RGG-rich NIQI motifs in proteins identified by RBPmap

