## Supplementary Material 3: Categorization of IRs according to their overlap with a feature or feature neighborhood for "In-depth Bioinformatic Analyses of Human SARS-CoV-2, SARS-CoV, MERS-CoV, and Other *Nidovirales* Suggest Important Roles of Noncanonical Nucleic Acid Structures in Their Lifecycles"

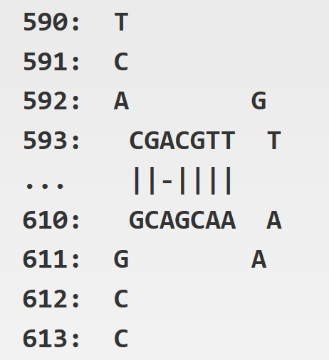

Supplementary Figure S1: Visualisation of inverted repeat. This IR has following parameters: size 7, spacer size 4, mismatch 1 (7-4-1).

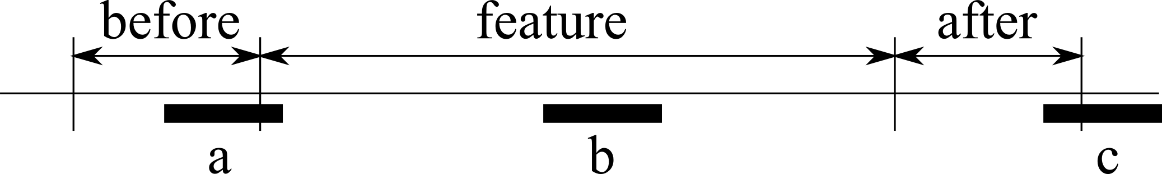

Supplementary Figure S2: Neighbourhood of an annotated feature. Example of possible IR occurrence around features and its classification: a) An inverted repeat is considered to be in near neighbourhood because it overlaps only partially with a feature. b) An inverted repeat overlapping fully with a feature and therefore is considered to be inside. c) An inverted repeat is not considered to be in near neighbourhood because it is not fully overlapping neither with a feature or its neighbourhood.
